# New Records of California Serogroup Virus in *Aedes* Mosquitoes and First Detection in Simulioidae Flies from Northern Canada and Alaska

**DOI:** 10.1101/2021.03.22.433603

**Authors:** Carol-Anne Villeneuve, Kayla J. Buhler, Mahmood Iranpour, Ellen Avard, Antonia Dibernardo, Heather Fenton, Cristina M. Hansen, Géraldine-G. Gouin, Lisa L. Loseto, Emily Jenkins, Robbin L. Lindsay, Isabelle Dusfour, Nicolas Lecomte, Patrick A. Leighton

## Abstract

An expected consequence of climate warming is an expansion of the geographic distribution of biting insects and associated arthropod-borne diseases (arboviruses). Emerging and reemerging arboviruses that can affect human health are likely to pose significant consequences for Northern communities where access to health resources is limited. In the North American Arctic, little is known about arboviruses. Thus, in 2019, we sampled biting insects in Nunavik (Kuujjuaq), Nunavut (Igloolik, Karrak Lake and Cambridge Bay), Northwest Territories (Igloolik and Yellowknife) and Alaska (Fairbanks). The main objective was to detect the presence of California serogroup viruses (CSGv) – a widespread group of arboviruses across North America and that is known to cause a wide range of symptoms, ranging from mild febrile illness to fatal encephalitis. Biting insects were captured twice daily for a 7-day period in mid-summer, using a standardized protocol consisting of 100 figure-eight movements of a sweep net. Captured specimens were separated by genus (mosquitoes) or by superfamily (other insects), and then grouped into pools of 75 by geographical locations. In total, 5079 *Aedes* mosquitoes and 1014 Simulioidae flies were caught. We report the detection of CSGv RNA in mosquitoes captured in Nunavut (Karrak Lake) and Nunavik (Kuujjuaq). We also report, for the first time in North America, the presence of CSGv RNA in Simulioidae flies. These results highlight the potential of biting insects for tracking any future emergence of arboviruses in the North, thereby providing key information for public health in Northern communities.

## Introduction

Rapid climate warming is altering ecological communities in the Arctic at an alarming rate (IPCC 2007; Post et al. 2009), thereby creating new habitats for biting insects and the diseases that they can carry (Bartlow et al. 2019). In spite of our increased awareness that the Arctic is experiencing climate warming twice as fast as the rest of the world (Jansen et al. 2020), studies documenting arthropod-borne viruses (arboviruses) in Northern regions are outdated (McLean et al. 1975, 1976, 1977a; McLean 1983), or in some cases, non-existent (i.e. vector species and arbovirus prevalence).

The most commonly reported arboviruses associated with biting insects in the Arctic are California serogroup viruses (CSGv; family *Bunyaviridae*, genus *Orthobunyavirus*) (Kurstak et al. 1979; Calisher 1996). They include the Snowshoe hare (SSH) and Jamestown Canyon (JC) viruses, which are occasionally associated with febrile and neuroinvasive disease in humans (LeDuc 1987). Their predominant arthropod vectors are mosquitoes of the genera *Aedes* and *Culiseta* (LeDuc 1987). For the SSH virus, the primary amplification hosts are snowshoe hares, squirrels and other small mammals, while for the JC virus, they are believed to be wild free-ranging ungulates (Drebot 2015). Even thought serological studies have shown that antibodies of both SSH and JC viruses are present in wildlife and people in the Arctic (Zarnke et al. 1983; Walters et al. 1999; Miernyk et al. 2019), there have only been a few reports documenting these viruses in mosquitoes captured in the field (McLean et al. 1975, 1977a, b).

There is less information available about other possible vectors for the CSGv, such as black flies (Simuliidae) and biting midges (Ceratopogonidae), both of which are present in Northern regions. Here, we report the results from our 2019 sampling effort aimed at detecting the presence of California serogroup viruses in Northern biting insects. For simplicity, the term *biting flies* will be used hereafter to describe biting insects that are not mosquitoes.

## Materials and methods

Insects were captured in the summer of 2019 in Northern Canada and Alaska (**Figure 1**). Locations were chosen based on a set of cities, villages or remote field sites identified by the Canadian Arctic One Health Network and the presence of local research partners willing to participate in surveillance activities. Sampling took place in Alaska (Fairbanks: 64.9152, −147.966), the Northwest Territories (Hendrickson Island: 69.8405, −133.975; Yellowknife: 62.5183, −114.320), Nunavut (Cambridge Bay: 62.1204, −105.045; Karrak Lake: 67.2359, −100.257; Igloolik: 69.3940, −81.3894) and the Nunavik region of Northern Québec (Kuujjuaq: 58.1272, −68.3848).

**Figure 1.**
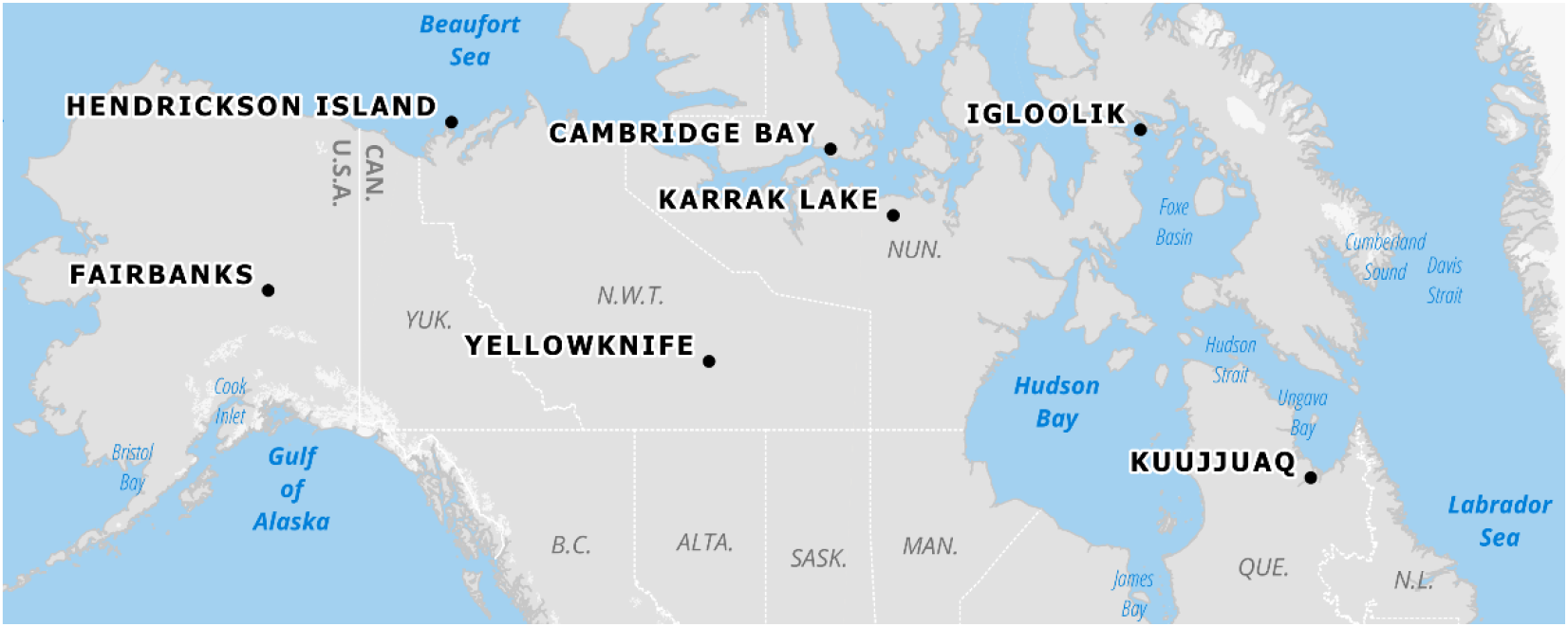
Sampling locations in 2019.

Manual capture, using a standardized protocol consisting of 100 figure-eight movements with an 18” sweep net (∼46 cm), was carried out twice daily (dawn and dusk) for 7 consecutive days in mid-summer of 2019. This technique is well-adapted to remote field site sampling where low equipment volume/weight and short sampling times are ideal (Silver 2008). It is also suitable for citizen scientists. Insects collected in the field were placed in a labeled plastic container and frozen at −18 °C until they were shipped to the Faculty of Veterinary Medicine, Université de Montréal, in Saint-Hyacinthe, QC.

All sorting of collected insects was conducted by a single observer (C.-A.V.) using a dissecting microscope and a chill tray. Mosquitoes and biting flies were separated from other insects. Subsequently, mosquitoes were sorted on a chill tray by genus using dichotomous keys (Wood et al. 1979; Thielman and Hunter 2004). Mosquitoes and biting flies were further grouped into pools by geographical location. Pools were stored at −80 °C until further analysis.

The pooled specimens were sent to the National Microbiology Laboratory in Winnipeg, MB for RNA extraction and real-time reverse transcriptase-polymerase chain reaction (RT-PCR) analysis. A sterile copper-coated steel bead (BB) and 1 ml of BA (made up of 10 ml of 10x M199 medium with Earls salts, 5 ml 1 M Tris buffer pH 7.6, 13.3 ml 7.5 % bovine serum albumin Fraction V and 1ml 100X penicillin/streptomycin) were added to each pool of mosquitoes and flies. Pools were homogenized using a TissueLyser (QIAGEN, Valencia, CA, USA) for 1 min at 30 Hz, and then centrifuged for 30 s at 13,000 rpm. RNA extraction was performed according to the manufacturer’s protocol for a RNeasy 96 Kit (QIAGEN). For each pool, 5 μl of eluted RNA was used for the RT-PCR reaction testing. The eluted RNA was combined with Applied Biosystems™ TaqMan™ Fast Virus 1-step master mix (ThermoFisher, Waltham, MA, USA). The target regions for the CSG, JC and SSH viruses were amplified using their respective primer pairs and probe (Wang et al. 2009) (**Table 1**). Positive controls were also used to test for CSG, JC and SSH viruses. Results were given as quantification cycle (Cq) units, i.e., the number at which the fluorescence of the probes increases, which is inversely related to the amount of RNA present (Williams 2009). Pools were considered positive when Cq units were ≤ 40.

**Table 1.**
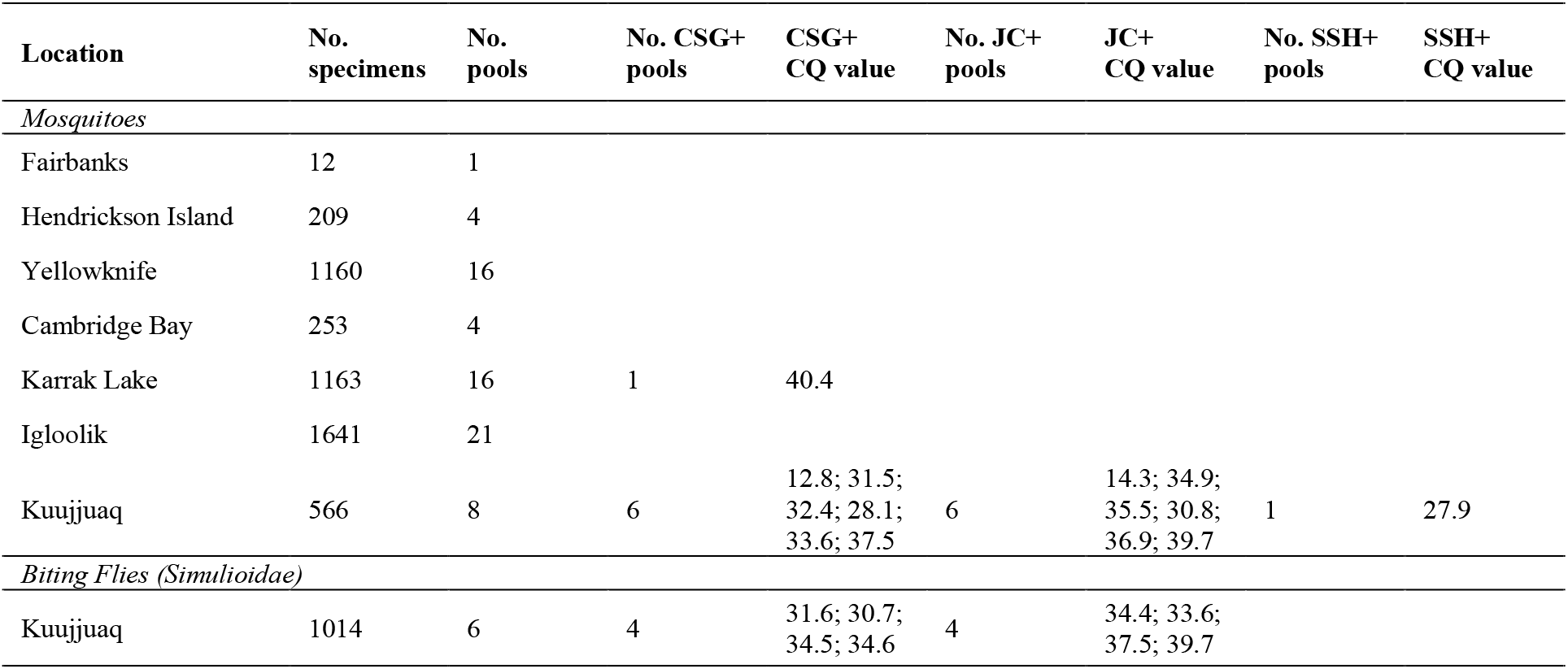
Mosquitoes and other biting flies collected from the North American Arctic, listed by geographical location from west to east (collected in mid-summer of 2019) and tested for CSG, JC and SHH viral RNA by RT-PCR

| Location | No. specimens | No. pools | No. CSG+ pools | CSG+ CQ value | No. JC+ pools | JC+ CQ value | No. SSH+ pools | SSH+ CQ value |
| --- | --- | --- | --- | --- | --- | --- | --- | --- |
| *Mosquitoes* |  |  |  |  |  |  |  |  |
| Fairbanks | 12 | 1 |  |  |  |  |  |  |
| Hendrickson Island | 209 | 4 |  |  |  |  |  |  |
| Yellowknife | 1160 | 16 |  |  |  |  |  |  |
| Cambridge Bay | 253 | 4 |  |  |  |  |  |  |
| Karrak Lake | 1163 | 16 | 1 | 40.4 |  |  |  |  |
| Igloolik | 1641 | 21 |  |  |  |  |  |  |
| Kuujuaq | 566 | 8 | 6 | 12.8; 31.5;<br>32.4; 28.1;<br>33.6; 37.5 | 6 | 14.3; 34.9;<br>35.5; 30.8;<br>36.9; 39.7 | 1 | 27.9 |
| *Biting Flies (Simuliidae)* |  |  |  |  |  |  |  |  |
| Kuujuaq | 1014 | 6 | 4 | 31.6; 30.7;<br>34.5; 34.6 | 4 | 34.4; 33.6;<br>37.5; 39.7 |  |  |

**Table 3.**
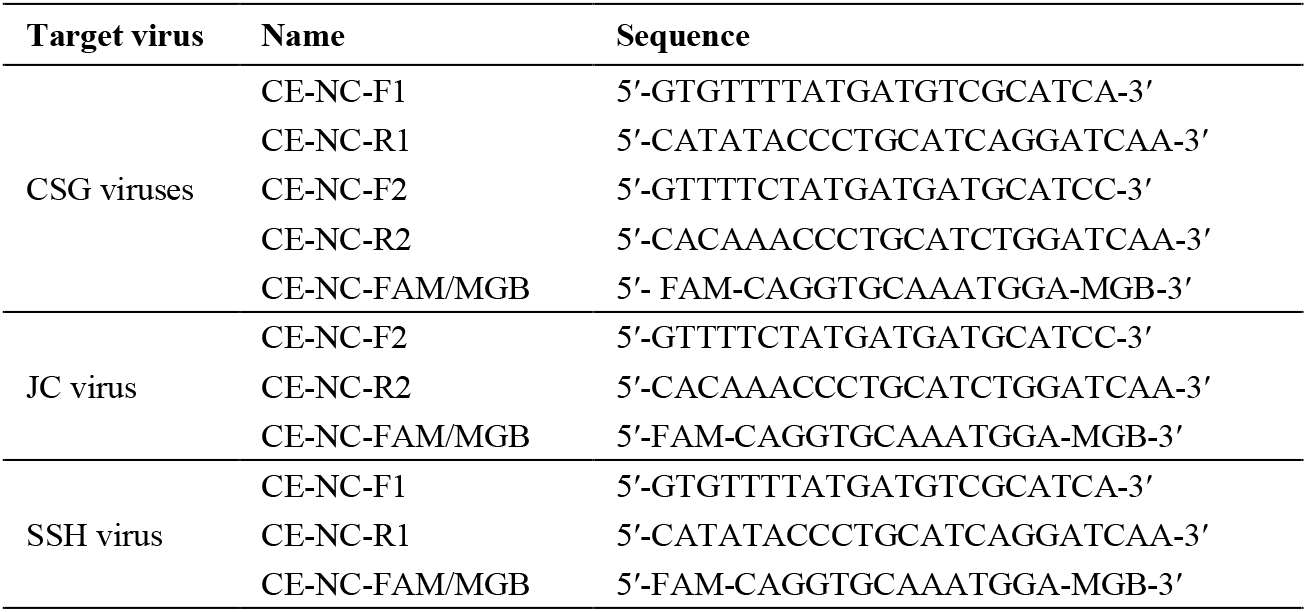
Probes and primer pairs for CSG, JC and SSH viruses

Positive pools were expressed as frequencies and 95% confidence interval calculated with R (R Core Team 2020) using *prop*.*test* (Newcombe 1998).

## Results

Between June 28^th^ and August 2^nd^ 2019, a total of 5079 mosquitoes and 1014 biting flies were caught (**Table 2**). Mosquitoes were collected across the seven sampling sites, forming 70 pools with an average of 75 specimens/pool (range: 12-86). Kuujjuaq was the only site where enough biting flies were collected to allow pooled analysis, forming 6 pools with an average of 169 specimens/pool (range: 114-237).

All collected mosquitoes were *Aedes* spp. females. This observation is consistent with our capture techniques, as sweep netting predominately captures females seeking a blood meal (Silver 2008). Biting flies appear to be predominantly black flies (Simulidae) because of their morphological characteristics: small, dark coloured with short legs, broad wings and a humpback appearance. A few smaller flies, probably biting midges, were also observed. Morphological observations were not made beyond the sorting process. Since biting flies were all pooled together, we decided to extend their identification to the superfamily level, Simulioidae, which consists of black flies and biting midges (Ceratopogonidae). For biting flies, 6 pools were made with an average of 169 specimens/pool (range: 114-237).

We tested each pool for CSG viruses, and then further tested CSG-positive pools specifically for JC and SSH viruses (**Table 2**). None of the 42 pools of mosquitoes collected in Fairbanks, Hendrickson Island, Yellowknife, Cambridge Bay or Igloolik tested positive for CSG viruses (**Table 2**). One of the 16 (6.2 %; 95 % CI [1.1, 28.3]) mosquito pools from Karrak Lake tested positive for CSG viruses. However, the quantity of viral RNA (cq of 40) in this sample was too low to further test for JC and SSH viruses with RT-PCR. For mosquitoes collected in Kuujjuaq, six of the eight (75 %; 95 % CI [40.9, 92.9]) pools tested positive for CSG viruses. Of these six pools, one (16.6 %; 95 % CI [3.0, 56.3] also tested positive for both JC and SSH viruses, while the five others (83.3 %; 95 % CI [43.7, 97.0] tested positive only for JC virus. We detected CSG viruses in biting flies in Kuujjuaq. Four of the six (66.6 %; 95 % CI [30.0, 90.3]) pools of biting flies tested positive for CSG viruses, which subsequently tested positive for JC virus.

## Discussion

In 2019, we detected CSGv RNA in mosquitoes at two of seven field sites across Northern Canada and Alaska. For the first time in North America, we also report CSGv in biting flies (Simulioidae).

All captured mosquitoes were from the genus *Aedes*, which is not surprising since *Aedes impiger, Aedes nigripes*, and *Aedes hexodontus* are the three main Arctic mosquito species (Ward and Darsie 2005). Moreover *Aedes* mosquitoes, like *Aedes communis*, are common vectors of CSGv, even in a sub-arctic environment (McLean et al. 1976, 1977b; Kurstak et al. 1979). However, species-specific results were not reported in this study, mainly because captured mosquitoes were too damaged for morphological identification past the genus level. There are alternatives to traditional identification methods, such as barcodes targeting the COI gene (Meier et al. 2006), but those methods have not yet been proven to be reliable for Arctic species. Reliable DNA databases from morphologically confirmed specimens are required before using these methods.

We detected CSGv (JC and SSH viruses) in mosquitoes captured in Nunavut (Karrak Lake) and Nunavik (Kuujjuaq). However, viral detection can be affected by multiple factors. For instance, temperature is known to influence vector, host, and arboviral distribution (Ciota and Keyel 2019). Since Kuujjuaq is the most southern and eastern site sampled (58.127277, −68.384854), its warmer summer could explain why positive pools were predominantly found there. Viral detection is also closely linked to species captured (Andreadis et al. 2008). For example, negative results in Hendrickson Island can simply mean that caught mosquitoes were not vectors for CSGv or that vectors were not active at the sampling time. More importantly, viral detection is highly dependant on pool size (Huang et al. 2001). For example, in Fairbanks, even if CSGv were present in one of the 17 mosquitoes caught, it might not have been detected due to low test sensitivity. Furthermore, 2019 results were not typical for Alaska, a state known for its abundance of mosquitoes. Without any baseline information about CSGv vectors in these regions, our ability to extrapolate results is therefore limited.

For the first time in North America, we detected CSGv (JC virus) in biting flies (Simulioidae) captured in Kuujjuaq. To the best of our knowledge, only one other study has detected CSGv in biting flies, which was in another biome, i.e., the former Czechoslovakia (Halouzka et al. 1991). Also, viral detection in biting flies (Simulioidae) does not necessarily indicate vector competence. It could simply indicate the ingestion of a viremic bloodmeal. Their role as potential vectors for needs further examination (Sick et al. 2019).

To conclude, results from this pilot year suggest that field collection of biting insects has the potential to provide useful data to monitor the changing distribution and local risk associated with vector-borne zoonoses across Northern regions. The sampling protocol used in this study provides a simple and cost-effective method for sampling biting insects in remote Northern communities, with potential for widespread application across the Arctic. Building reliable DNA databases for potential vectors as well as subsequent monitoring years will be useful to establish a baseline on current species’ ranges and infective status. Such studies are crucial to anticipate and track any future emergence of arboviruses in the North, thereby providing key information for public health in Northern communities.

## Acknowledgments

We would like to thank Mathieu Archambault (Université de Moncton), Ashley Elliott (Fisheries and Oceans Canada), Camille Guillot (Université de Montréal), Robin Owsiacki and Adrián Hernández Ortiz (University of Saskatchewan), Megan Wardekker (Fisheries and Oceans Canada), Jordan Bertagnolli (Government of the Northwest Territories) and the Centre for Biodiversity Genomics (University of Guelph) for their help in collecting mosquitoes in the field. We would like to thank Scott G. Harroun (Université de Montréal) for his help in editing this manuscript.

We would also like to thank Patricia Lacroix (Government of the Northwest Territories) and Malik Awan (Government of Nunavut) for their help with our Wildlife Research Permit applications, as well as all of the organizations and communities who supported our project: Olokhaktomiut Hunter’s And Trapper’s Committee, Tuktoyaktuk Hunters and Trappers Committee, Wildlife Management Advisory Council (Northwest Territories), Environmental Impact Screening Committee (Inuvialuit Settlement Region), North Slave Metis Alliance, Aklavik Hunters and Trappers committee, Inuvik Hunters & Trappers, the Tłįcho Government Department of Culture and Lands Protection as well as the Hunter and Trapper Organisations of Igloolik.

This study was carried out through the Canadian Arctic One Health Network (CAOHN), with funding from ArcticNet (Networks of Centres of Excellence of Canada), Polar Knowledge Canada (POLAR), and the Natural Sciences and Engineering Research Council of Canada (NSERC)

